# Cross-feeding between *Thauera aminoaromatica* and *Rhodococcus pyridinivorans* drove quinoline biodegradation in a denitrifying bioreactor

**DOI:** 10.1101/2020.01.31.929745

**Authors:** Xinxin Wu, Xiaogang Wu, Ji Li, Qiaoyu Wu, Yiming Ma, Weikang Sui, Liping Zhao, Xiaojun Zhang

**Author notes:** Corresponding author: Dr. Xiaojun Zhang, 800 Dongchuan Road, Minhang District, Shanghai, 200240, China.

## Abstract

The complex bacterial community is predominated by several taxa, such as *Thauera* and *Rhodococcus*, in a quinoline-degrading denitrifying bioreactor. Yet it remains unclear about how the interactions between the different bacteria mediate the quinoline metabolism in denitrifying condition. In this study, we designed a sequence-specific amplification to guide the isolation of the most predominant bacteria and obtained four strains of *Thauera aminoaromatica,* the representative of one key member in the bioreactor. Test on these isolates demonstrated that all of them were unable to strive on quinoline but could efficiently degrade 2-hydroxyquinoline, the hypothesized primary intermediate of quinoline catabolism, under nitrate-reducing condition. However, another isolate, *Rhodococcus pyridinivorans* YF3, corresponding to the second abundant taxon in the same bioreactor, was found to degrade quinoline via 2-hydroxyquinoline. The end products and removal rate of quinoline by isolate YF3 were largely varied with the quantity of available oxygen. Specifically, quinoline could only be converted into 2-hydroxyquinoline without further transformation under the condition with insufficient oxygen, e.g. less than 0.5% initial oxygen in the vials. However, if were aerobically pre-cultured in the medium with quinoline the resting cells of YF3 could anaerobically convert quinoline into 2-hydroxyquinoline. A two-strain consortium constructed with isolates from *Thauera* (R2) and *Rhodococcus* (YF3) demonstrated an efficient denitrifying degradation of quinoline. Thus, we experimentally proved that the metabolism interaction based on the 2-hydroxyquinoline cross-feeding between two predominant bacteria constituted the mainstream of quinoline degradation. This work sheds light on the understanding of mechanism of quinoline removal in the denitrifying bioreactor.

**Importance:** We experimentally verified the most predominant *Thauera* sp. was indeed active degrader for the intermediate metabolites and the second abundant taxon *Rhodococcus* exerted, however, key function for opening the food box for a complex quinoline-degrading community. An ecological guild composed of two isolates was assembled, revealing the different roles of keystone organisms in the microbial community. This study, to our best knowledge, is the first report on the cross feeding between the initial attacker with unprofitable catalysis of reluctant heterocyclic compounds and the second bacterium which then completely degrade the compound transformed by the first bacterium. These results could be a significant step forward towards elucidation of microbial mechanism for quinoline denitrifying degradation.

## Introduction

Quinoline and its derivatives are typical N-heterocyclic compounds that occur widely in coal tar, shale oil and creosote, and serve as raw materials and solvents in chemical, pharmaceutical and pesticide industries (1). They are known to be carcinogenic and mutagenic to human (2,3) and aroused a significant concern as recalcitrant pollutants to ecological environment.

Anaerobic bioremediation is an attractive technology due to its virtue of energy saving and cost-effectiveness, since heavily contaminated environments are often oxygen deficient (4). However, most of the literatures focused the aerobic degradation of quinoline. Various microorganisms capable of metabolizing quinoline aerobically have been isolated, mostly belonging to *Pseudomonas* (5–7), *Rhodococcus* (8) and *Bacillus* (9). The pathways of aerobic quinoline degradation have also been well described (10). However, little attention was paid to the anaerobic quinoline biodegradation. Degradation of quinoline in industrial scale wastewater treatment reactors had been reported in a few literature (11,12). In addition, no evidence proved the role of main anaerobic degraders in these industrial bioreactors. There were several efforts to elucidate the anaerobic degraders by using the lab scale bioreactor and batch culture experiments (13–16). But to date, only one isolate, *Desulfobacterium indolicum* strain DSM 3383, which used sulfate as electron acceptor, was purely cultured as an anaerobic quinoline degrader (17). It is wrapped in mystery why there is few isolate for anaerobic quinoline degradation.

An efficient anoxic microbial community was enriched in a chemostat that was operated for more than 10 years with quinoline as electron donor and nitrate as electron acceptor. Phylogenetic analysis of this consortium showed that specific phylotypes were associated with different stages of the degradation (16), which suggests that microorganisms interact during quinoline metabolism. However, our understanding of this interaction is restricted due to the complexity of the community composition in the bioreactor while lack of anaerobic degrading microorganisms available in pure culture. Therefore, functional analysis using the representative isolates would be a crucial further step for understanding the quinoline denitrifying degradation in the reactor.

To investigate the underlying microbial processes in this complex quinoline-degrading consortium, we endeavored to isolate the most predominant and active bacteria, which had been identified as the keystone organisms involved in denitrifying quinoline removal (16,18). The degradation characteristics of these isolates were evaluated under different conditions. Based on the degradation function of different isolates, a co-culture of representative isolates of *Thauera* and *Rhodococcus* were constructed to demonstrate the cooperation of two bacteria during quinoline metabolism under defined conditions.

## Materials and methods

### Operation of the quinoline denitrifying bioreactor

A 2-L tank, filled with plastic rings and synthetic fiber strings as semisoft media, was used to construct a lab-scale bioreactor. Seeding sludge was collected from an anoxic tank of a coking wastewater treatment plant from the Shanghai Coking and Chemical Factory (Wujing, Shanghai). The synthetic wastewater was composed of quinoline (100 mg/L), NaNO_3_ (240 mg/L) and K_2_HPO_4_ (140 mg/L). During wastewater upflowing (i.e. from the bottom to up) without any agitation, the inner of reactor was oxygen-depleted due to the rapid consumption by aerobic bacteria. The hydraulic retention time was 24 h. The pH and temperature of the reactor were adjusted and controlled at 7.5 and 25 °C, respectively.

### Profiling of the quinoline-degrading bacterial community

The biofilm sample was collected by scraping the biofilm from the surface of the supporting materials in the bioreactor. Genomic DNA extraction of the samples in triplicate was conducted as previously described (19). The sequencing library of V3-V4 regions in the 16S rRNA gene were constructed by two-step PCR amplification according to Illumina’s instructions. The purified amplicons were sequenced using the Illumina MiSeq System (Illumina Inc., United States). The preliminary process of the raw sequencing data was conducted as the previous document (20). Sequence assembly was first implemented, and the unique sequences obtained by dereplication were sorted by decreasing abundance, and then singletons were abandoned. UPARSE’s default (21) was used to select the representative operational taxonomy units (OTUs), and UCHIME (22) was selected to further perform reference-based chimera detection against the RDP classifier training database (23). Finally, the OTU table was completed by mapping quality-filtered reads to the representative OTUs with Usearch (24), resulting in a global alignment algorithm at a 97% cutoff. Further analysis was performed using the QIIME platform (version 1.8)(25). In addition, representative sequences for each OTU were submitted to the online RDP classifier (RDP database version 2.11) to determine the phylogeny, with a bootstrap cutoff of 80%. The 16S rRNA gene sequences in this study were submitted to the GenBank Sequence Read Archive (SRA) database in the National Center for Biotechnology Information (NCBI) under the accession number SRP188486.

### Isolation and identification of the most predominant bacteria

Six types of media were used to isolate the most predominant *Thauera* spp. strains. The composition of the six media were described as follows, Nutrient Agar (NA): peptone 10 g/L, beef extract 3 g/L, NaCl 5 g/L, agar 15 g/L (26); 1/10 strength NA: diluted by 10 folds based on NA except for the content of agar; Tryptic Soy Agar (TSA): tryptone 17 g/L, soy peptone 3 g/L, glucose 2.5 g/L, NaCl 5 g/L, K_2_HPO_4_ 2.5 g/L, agar 15 g/L (27); 1/10 strength TSA: diluted by 10 folds based on TSA except for the content of agar; Reasoner’s 2A agar (R2A): yeast extract 0.5 g/L, peptone 0.5 g/L, casein hydrolysate 0.5 g/L, glucose 0.5 g/L, soluble starch 0.5 g/L, KH_2_PO_4_ 0.3 g/L, MgSO_4_ 0.024 g/L, sodium pyruvate 0.3 g/L, agar 15 g/L (27); Waste Water Medium (WWM): made by raw wastewater which was filtrated and removed bacteria and supplemented with 1.5% agar.

The specific PCR primer targeted the most predominant *Thauera* sp. was designed by DNAMAN (version 7.0) using the sequences of different OTUs belonging to genus *Thauera,* which obtained from a full length 16S rRNA clone library constructed for quinoline denitrifying bioreactor (18). OTU specific primers were used to assist the screening of the target *Thauera* strains following the previously described procedure (13).

The biofilm sample was obtained and mixed in a shaker with 150 rpm for 2 h. Then the suspension was diluted and spread on the plate of six types of media including NA, 1/10 strength NA, TSA, 1/10 strength TSA, R_2_A and WWM. All plates were incubated at 30 °C under aerobic and anaerobic conditions, respectively. The isolates that had positive signal of colony PCR were purified by plate-streaking technology. Finally, genomic DNA of all isolates were extracted by phenol-chloroform protocol (28). The ERIC-PCR was conducted to have genomic typing of all isolates (29). And representative strains for different type of ERIC fingerprints were selected for 16S rRNA gene sequencing to identify the taxonomy and construct phylogenetic tree. The 16S rRNA gene sequences of representative strains were deposited in GenBank with the accession numbers of MK271352 (R2), MK272943 (R4), MK272944 (N15), MK272945 (N38), respectively.

### Pre-culture of bacterial inoculum for biodegradation experiment

Unless specific mention, the inoculum for all experiments was prepared by inoculating the strains of *Thauera* spp. or *Rhodococcus pyridinivorans* YF3 (GU143680.1) in the Nutrient Broth medium (NB) and incubating at 30 °C, and 150 rpm on a rotary shaker for 24 h. The bacterial cells were harvested by centrifugation and washed three times in mineral salt medium (MSM) and the suspension was used as inoculum. The mineral salt medium (MSM) contained: K_2_HPO_4_·3H_2_O 0.57 g/L, KH_2_PO_4_ 0.233 g/L, NH_4_Cl 0.02675 g/L, NaCl 0.5 g/L, NaNO_3_ 85 mg/L (1mM) and trace elements including NaHCO_3_ 0.168 g/L, MgSO_4_ 0.12 g/L, CaCl_2_ 0.0544 g/L, disodium EDTA 0.025 g/L, H_3_BO_3_ 0.0036 g/L, FeSO_4_·7H_2_O 0.0015 g/L, CoCl_2_ 0.0012 g/L, Ni(NH_4_)_2_(SO_4_)_2_ 0.0012 g/L, Na_2_MoO_4_ 0.00094 g/L, Na_2_SeO_4_ 0.00026 g/L, MnSO_4_ 0.0002 g/L, ZnSO_4_ 0.00016 g/L, CuSO_4_ 0.000032 g/L, pH 7.4.

### Degradation experiments for bacterial isolates

The batch experiments were conducted using a series of 20 mL headspace vials, which contained 10 mL of either the MMQ or MMHQ medium. The composition of the MMQ or MMHQ were described as follows. MMQ: mineral salt medium (MSM) supplemented with 0.23 mM Quinoline (Sigma–Aldrich, Co., Inc); MMHQ: mineral salt medium (MSM) supplemented with 0.23 mM 2-hydroxyquinoline (Sigma-Aldrich, Co., Inc).

The inoculum was 5% to 10% of volumetric proportion. For each of the different media, vials without inoculum but maintained under the same condition were set as the abiotic negative control. All vials were immediately sealed with airtight butyl-rubber septa and aluminum crimp caps and made anoxic by repeated evacuation and filling with helium. To investigate the effects of pH value on the degradation, the initial pH of media was adjusted to 4.5, 6.5, 7.5, 8.5 and 10.5, respectively.

To investigate the effects of oxygen on quinoline biotransformation by *Rhodococcus* sp. YF3, 0.04 mL, 0.10 mL and 0.16 mL pure oxygen were respectively injected into the vials by precision syringes, named as the groups of 0.2% O_2_, 0.5% O_2_, 0.8% O_2_, respectively. The headspace of anaerobic vials was no oxygen injection while the headspace of aerobic vials was replaced by air. All vials were incubated in the dark at 30 °C with shaking at 150 rpm. Samples were taken periodically by syringes in anaerobic workstation to prevent from oxygen during incubation. All treatments in the above degradation experiment had triplicates to minimize the experimental variation.

### Measurement of chemical compounds

The sampled liquid was centrifuged and the supernatant was used for further analysis. The concentration of quinoline and 2-hydroxyquinoline were analyzed by HPLC system (Agilent, ZORBAX SB-C18 reverse-phase column, 150 × 4.6 mm, 5μm). The mobile phase was methanol solution with a volume ratio of 60:40 (methanol / water) at a flow rate of 1.0 mL/min. Quinoline and 2-hydroxyquinoline were both detected at 225 nm wavelength. The injection volume was 20 μL and column temperature was at 30 °C. The nitrate concentration was measured using a PXJ-1B ion meter (Jiangfen, China) with a pNO_3_ ^−1^ nitrate ion-selective electrode (Tianci, Shanghai).

## Results

### Profiling of the quinoline-degrading bacterial community

The removal rates of quinoline and nitrate maintained stably at about 80% and 90%, respectively, in the denitrifying quinoline-degrading bioreactor. Before sample collection for isolating the most predominant organisms, the bacterial community of triplicate samples from the reactor was profiled by high-throughput sequencing. The Illumina Miseq platform yielded 73902 high-quality reads with an average of 24634 reads per sample (±1418 SD). 460 species-level operational taxonomic units (OTUs) were delineated using 97% identity as cut-off value. Six predominant OTUs contributed to 52.06% of all reads (Fig. 1). The most predominant OTU (OTU1) with relative abundance of 22.5% belonged to genus *Thauera*; while OTU2, the second abundant OTU (12.5%), was affiliated to genus *Rhodococcus*.

**Figure 1.**
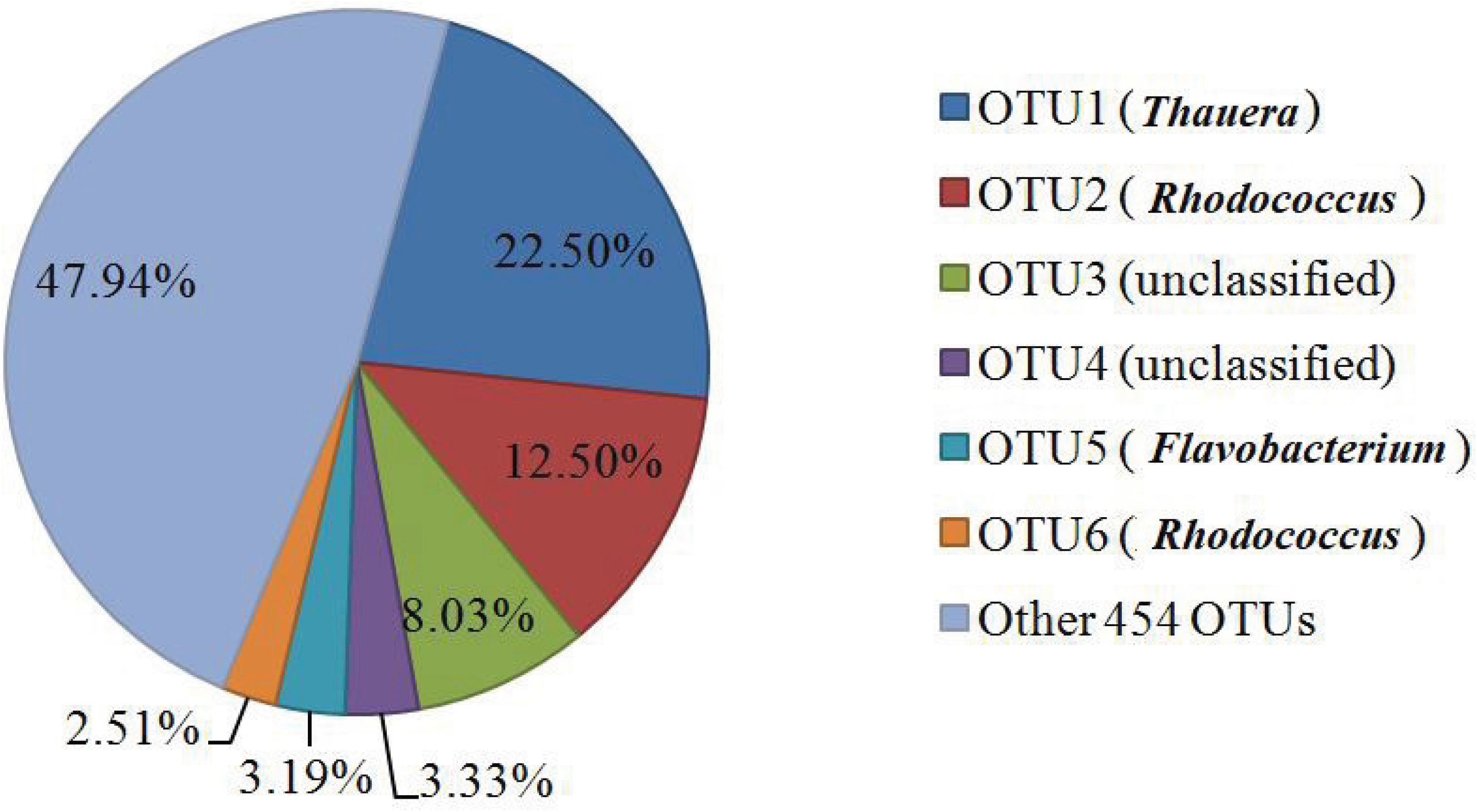
The abundance of six most predominant OTUs among 460 OTUs from the quinoline-degrading bioreactor.

### Isolation and identification of the most predominant bacteria

Since that the most predominant OTU was supposed as the key player for the quinoline degradation, we attempted to isolate the corresponding bacteria from the bioreactor using sequence-guiding isolation strategy. The specific PCR primers based on most predominant *Thauera* sp. were designed in silico. After selection based on the amplification, the primer pair of Thau135/Thau716 (5’-GGGATAACGTAGCGAAAGCT-3’; 5’-CCATCGGTGTTCCTCCTG-3’) was further evaluated experimentally for its specificity. The amplicon of reactor sample by using the selected primer pair was cloned and 15 clones were randomly selected for sequencing of partial 16S rRNA gene with length longer than 500 bp. Among them, 8 clones showed 100% identity with DR-80 and most predominant OTU1, and the other 5 clones shared over 99.7% identity with DR-80/OTU1, only 2 clones showed identity about 99.5%. These results indicated that the specificity of Thau135/Thau716 was high enough to guide the following isolation.

Several hundreds of colonies grown on the plates of six different media were screened by specific amplification using the primers of Thau135/Thau716. Totally, 13 positive colonies were picked, of which 3 were purified from R2A medium, 6 from 1/10 strength TSA and 4 from 1/10 strength NA medium. Morphologically, these selected colonies were pearl-like on the 1/10 NA plate and the cells formed characteristic flocs or clusters in NB liquid culture shaking with 150 rpm at 30 °C. All of these isolates were classified into four genotypes by ERIC-PCR result, in which the isolates of R2, R4, N15 and N38 were selected as the representative strains for each ERIC type. The full length of 16S rRNA gene sequences of four representative strains were 100% identity with each other and had one nucleotide mismatch with the uncultured *Thauera* clone DR-80, which had highest abundance in a lab scale bioreactor for quinoline degradation (18). Besides, all isolates showed 100% identity with the 16S rRNA gene sequence of the most predominant *Thauera* genome which was assembled using the metagenome data for the same bioreactor in our previous work (30). Moreover, *Thauera aminoaromatica* strain S2 was the closest neighbor of all isolates when blast 16S rRNA gene sequences with NCBI nr database (Fig. 2), which was consistent with the previous report about the metagenome data that the assembled genome of the most predominant organism in the reactor was most closely related to *Thauera aminoaromatica* S2 (30). These four isolates phylogenetically discriminated with other three isolates, 3-35, Q4, Q20-C, which were obtained previously from a quinoline containing coking wastewater denitrifying treatment bioreactor from which the seeding sludge for our bioreactor was obtained (14).

**Figure 2.**
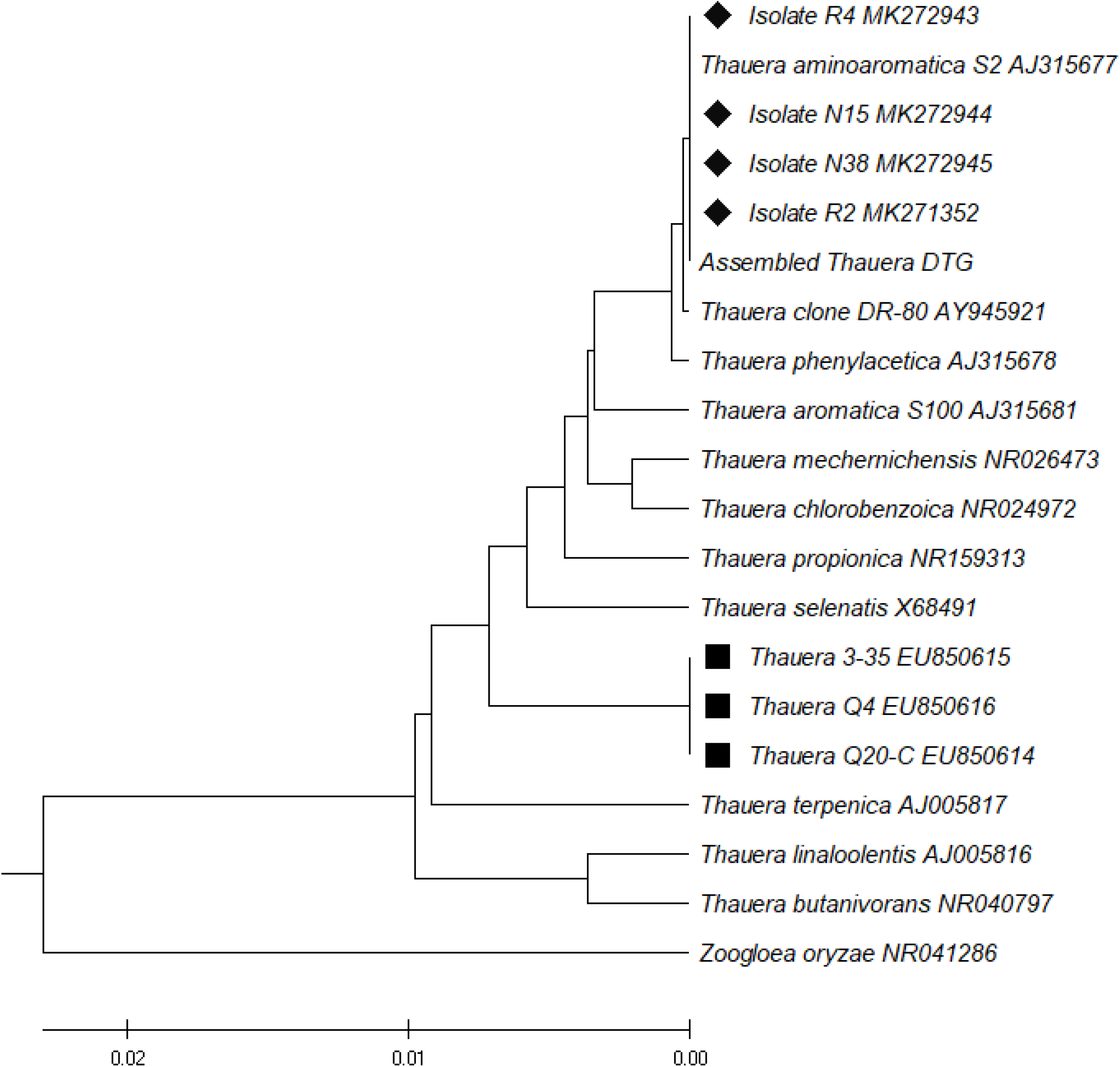
Phylogenetic tree based on the 16S rRNA gene sequences of isolates from quinoline degrading bioreactor and other related *Thauera* species from NCBI database. Numbers in bracket are the sequences accession number in GenBank. The bootstrap values of Neighbor-Joining analysis were labeled at the nodes. Diamond label means isolates in this study; square label means isolates from same reactor in previous study.

### Metabolism of Quinoline and 2-hydroxyquinoline by Thauera isolates

Firstly, the isolated *Thauera* strains were tested for their capacity of quinoline degradation. Unexpectedly, none of them could metabolize quinoline, when it was used as sole carbon source, both under anaerobic and aerobic conditions. However, all four isolates could utilize 2-hydroxyquinoline, the derivative of quinoline, which was reported as the first degrading intermediate of quinoline (31). Data showed that about 80% of 50 mg/L 2-hydroxyquinoline was eliminated in MMHQ medium within 10 days under nitrate-reducing condition and no any accumulation of intermediates in the measured samples (Fig. 3a). The effects of pH value on 2-hydroxyquinoline degradation were further investigated. The results showed that 30 mg/L 2-hydroxyquinoline was depleted in 132 h both under pH 7.5 and 8.5 conditions (Fig. 3b). Degradation of substrates was apparently inhibited under acidic (pH≤6.5) or alkaline (pH≥10.5) condition. The transformation of 2-hydroxyquinoline was pH-dependent, with the optimal pH range from 7.5 to 8.5. Additionally, although all isolates had capacity for 2-hydroxyquinoline removal, their rates for degradation were different. For example, the isolate N38 had the lowest rate, whereas the isolate R2 had the highest rate of 0.583 mg/L.h.

**Figure 3.**
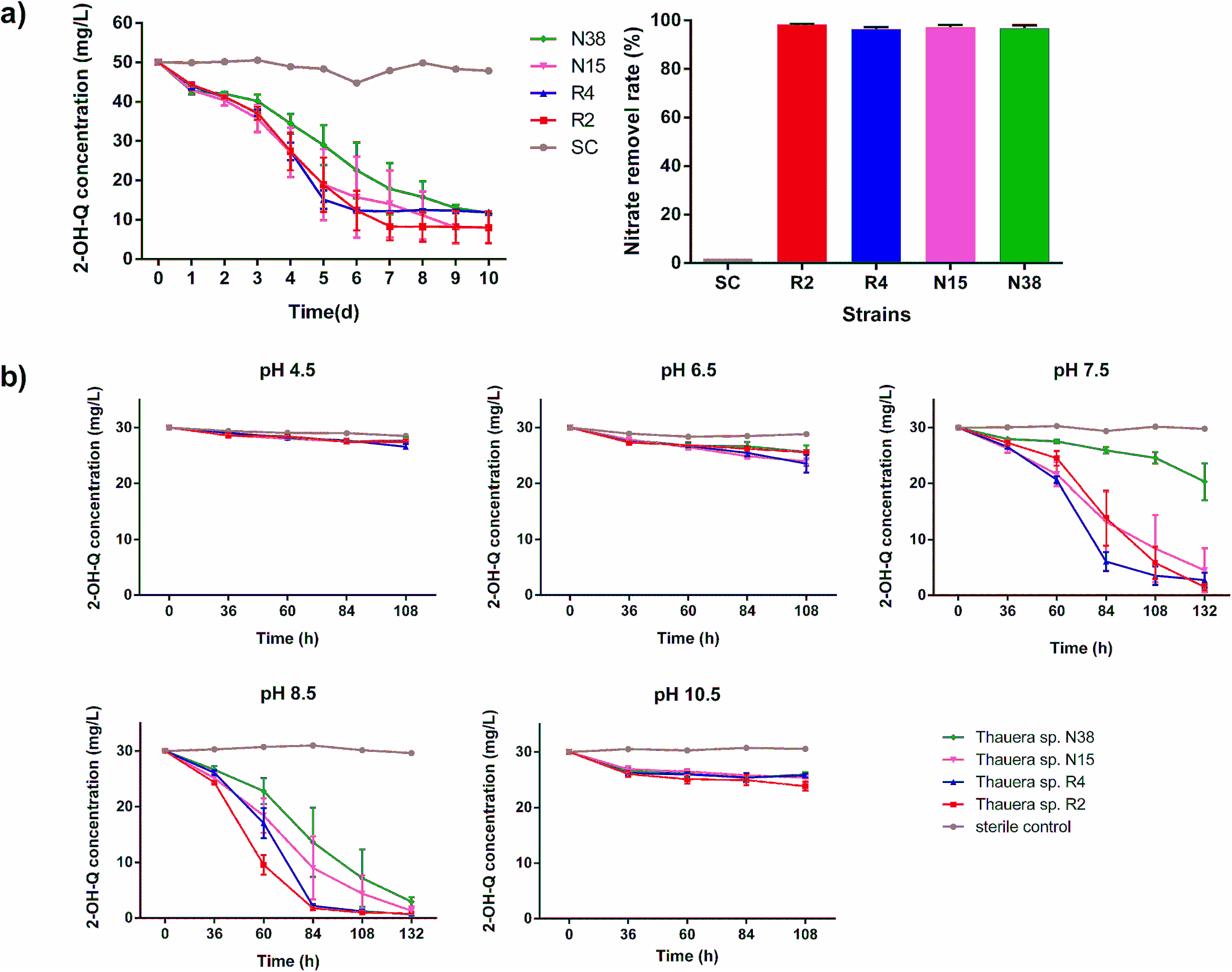
Dynamics of 2-hydroxyquinoline (2-OH-Q) and nitrate removal by four *Thauera* strains. a) degradation of 50 mg/L 2-OH-Q and 1mM nitrate degradation; b) the performance of 30 mg/L 2-OH-Q degradation under different pH conditions. SC represents the sterile control.

### Metabolism of quinoline by a *Rhodococcus* isolate

On the petri dish for bacteria isolation, a type of reddish colony frequently appeared. The identification by 16S rRNA gene suggested it represented another predominant bacterial phylotype for quinoline degradation proposed in previous literature (16). The isolate YF3 was selected as the representative of the most predominant *Rhodococcus* OTU, which was identified as *Rhodococcus pyridinivorans*. Transformation of quinoline by YF3 under different amount of initial oxygen in headspace of vials was explored. We found that the quinoline removal rate and transformed products were varied due to the different condition of oxygen supplement. After 24 h incubation, the degradation of microcosms which inoculated with the inoculum grown on NB medium was analyzed. The results showed that 0.23 mM quinoline was completely oxidized when supplemented with sufficient oxygen; in contrast, quinoline was persistent under anaerobic condition; whereas, about 0.09 mM 2-hydroxyquinoline accumulated and 0.13 mM quinoline remained if 0.2% oxygen supplied, which equal to 2.8 mg/L dissolved oxygen; quinoline was completely transformed into 2-hydroxyquinoline under 0.5% oxygen,, which equal to 7.1 mg/L dissolved oxygen; when the oxygen concentration increased to 0.8%, which equal to 11.2 mg/L dissolved oxygen, only 0.17 mM 2-hydroxyquinoline retained due to the further degradation (Fig. 4).

**Figure 4.**
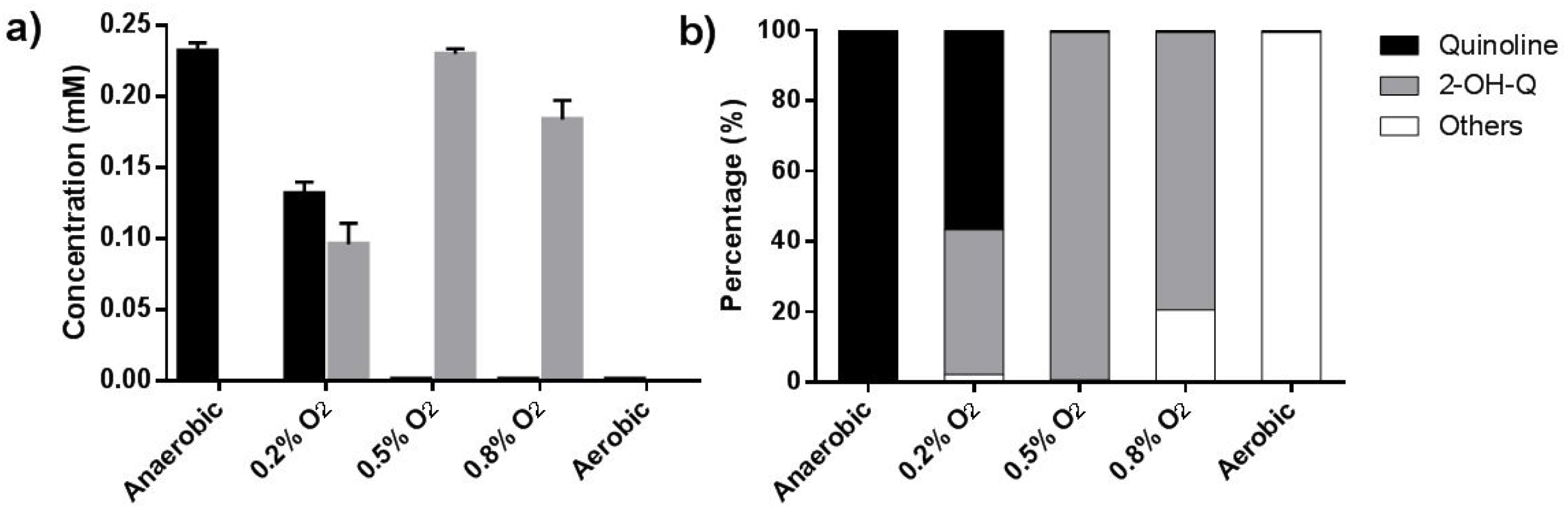
Transformation of quinoline by *Rhodococcus pyridinivorans* YF3 under different oxygen availability a) Concentration of quinoline and 2-hydroxyquinoline (2-OH-Q); b) The proportion of different components remained in cultures.

### Synergistic degradation of quinoline by a bacterial co-culture

Since that *Thauera* sp. R2 has been proved as an anaerobic 2-hydroxyquinoline degrader and *Rhodococcus* sp. YF3 had the capacity for hydroxylation of quinoline, we assumed that the degradation of quinoline might be accomplished in the bioreactor via metabolic cooperation of these two bacteria. Thus, we performed a trial of co-cultivation using the above mentioned bacterial strains of *Thauera* and *Rhodococcus* to test the capacity of degradation of quinoline under the conditions of different oxygen-availability. We found that 0.07 mM and 0.01 mM quinoline remained in the vials of experimental groups of 0.2% O_2_ and 0.5% O_2_, respectively. 2-hydroxyquinoline was finally eliminated in all the groups within 48 h while quinoline was completely depleted only in 0.8% O_2_ (Fig. 5a).

**Figure 5.**
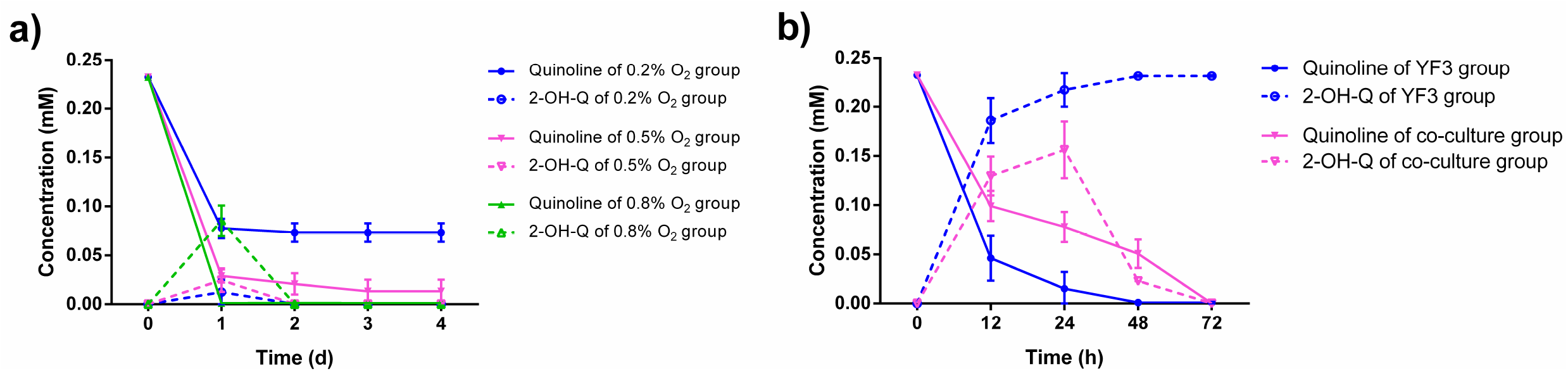
Dynamics of quinoline degradation by the co-culture of *Rhodococcus* sp. YF3 and *Thauera* sp. R2. a) under limited oxygen supplement with the inoculum cultured in NB medium; different colors represent different oxygen supplied conditions. b) under anaerobic condition with the inoculum precultured aerobically in medium containning quinoline. Blue line represents *Rhodococcus* sp. YF3 alone, while red line represents co-culture of *Rhodococcus* sp. YF3 and *Thauera* sp. R2. Solid line in a) and b) represents the change of quinoline concentration while the dotted line represent the change of 2-hydroxyquinoline (2-OH-Q) concentration.

Considering the original bioreactor is still an oxygen deficient system, we try to elucidate the possible mechanism of anaerobic quinoline degradation. We have verified that *Rhodococcus* sp. YF3 could hydroxylate quinoline under micro-oxygen condition. Washing and transferring YF3 cells, which were pre-incubated with quinoline, 2-hydroxyquinoline or phenol for 24 h under 0.5% oxygen condition, to fresh MMQ medium and incubating for another 72 h under completely anaerobic condition, we observed that 0.23 mM quinoline was basically transformed into equivalent 2-hydroxyquinoline by YF3 monoculture. 2-Hydroxyquinoline was further removed in the co-culture of YF3 and R2 (Fig. 5b).

## Discussion

In this study, we aimed to elucidate the microbial mechanism of quinoline denitrifying degradation. As a first step, we tried to isolate the most predominant bacteria in the quinoline-degrading consortium, which was considered as the promising anaerobic quinoline degrader. However, many attempts failed at isolating the target strain by using the media with quinoline as sole carbon source. In the earlier study, a *Thauera* genus-specific nested-PCR denaturing gradient gel electrophoresis (DGGE) method was developed, combining with media containing various carbon source, to guide the isolation of *Thauera* spp.. In that study, three *Thauera* strains (Q4, Q20-C, and 3-35) were obtained but none of them could utilize quinoline either aerobically or anaerobically (13,14). Considering that the former isolates were corresponding to the less abundant *Thauera* species in the bioreactor, in this study, we designed OTU specific PCR primers targeting the most predominant *Thauera* OTU1, as the biomarker to guide the isolation of the target *Thauera* bacterial strains. The colonies grown on various media were screened by using this method. As a result, four isolates belonged to *Thauera aminoaromatica*, were identified as the homologue of the most predominant bacteria in quinoline degrading bioreactor. *Thauera aminoaromatica* had been reported as one of denitrifying species capable of growing with amino-aromatic compounds (32). In this study, all isolates belonging to *Thauera aminoaromatica* were able to degrade 2-hydroxyquinoline but not quinoline under denitrifying conditions, which exactly explained previous unsuccessful attempts via the strategy of quinoline-degrading function-guiding isolation by using quinoline as the solo carbon source. Hence, sequence-guiding approach to assist isolating some difficult-to-culture bacteria could be an alternative and promising method.

Dozens literatures have reported that isolated bacteria may lack of the capability of complete mineralization of hydrocarbon, especially for the recalcitrant compounds (33). Moreover, regardless of present or absent of oxygen, the first metabolite of quinoline was 2-hydroxyquinoline (10). This compound was thus preferentially considered as substrate of most predominant *Thauera* bacteria in the denitrifying bioreactor. The results of 2-hydroxyquinoline degradation by monoculture of *Thauera* isolates in this study proved our hypothesis.

To further identify the bacteria being responsible for transforming quinoline to 2-hydroxyquinoline, we further explored other key players from the predominant bacteria in the community. The members of genus *Rhodococcus* were known as aerobic quinoline degraders (8), and this phylotype has been reported closely related with quinoline denitrifying degradation (15, 16). Based on that, *Rhodococcus pyridinivorans* YF3 isolated from the bioreactor was considered as a candidate to initial attack for quinoline in the denitrifying bioreactor. Unsurprisingly, the experimental result was well in accord with our speculation. Notably, the consumption of quinoline by strain YF3 depended upon the oxygen concentrations. Therefore, long-term accessing to low oxygen likely shape a *Rhodococcus*-dominated community in the quinoline denitrifying bioreactor (15).

Communities of bacteria are extraordinarily complex with hundreds of interacting taxa but the knowledge about how the tangled interactions within natural bacterial communities mediate ecosystem functioning is limited (34). A promising way to surpass this limitation is to create a synthetic community by artificially co-culturing of selected two or more species under a well-defined media (35). Considering that the metabolite of quinoline transformed by isolate YF3 could be further metabolized by isolate R2, we constructed a synthetic consortium using pure cultures of these two isolates to achieve cooperative degradation of quinoline. In this consortium, *Thauera* sp. R2 cross-feed on the 2-hydroxyquinoline released by *Rhodococcus* sp. YF3. Ecologically, this type of microbial interaction is equivalent to a commensalism (36). Here the initial step of quinoline degradation is hydroxylation at the 2 position which requires energy (37), but not provides any substrate and energy for growth. Accordingly, *Rhodococcus* sp. YF3 would not benefit from the hydroxylation reaction. In this study, the product 2-hydroxyquinoline of the first transformation step served as the substrate for the next organism in the chain. Meanwhile, *Thauera* sp. R2 released some intermediates during further metabolism. And the bacteria of *Rhodococcus* sp. YF3 could utilize these metabolites leaked from *Thauera* cells to obtain energy and for growth. Consequently, most of the carbon from quinoline was consumed by R2 cells and small portion by YF3 and other bacteria. It was well consistent with the highest abundance of *Thauera* sp. R2 and relatively lower abundance of *Rhodococcus* sp. YF3 in the bioreactor, although the latter acted like a lord to open the food box for *Thauera* R2. Thus, the cross-feeding and interdependence between these two strains was main driven force for quinoline denitrifying degradation and assembly for a stable community in the denitrifying bioreactor. We summarized the ecological mechanism of quinoline denitrifying degradation in Figure 6.

**Figure 6.**
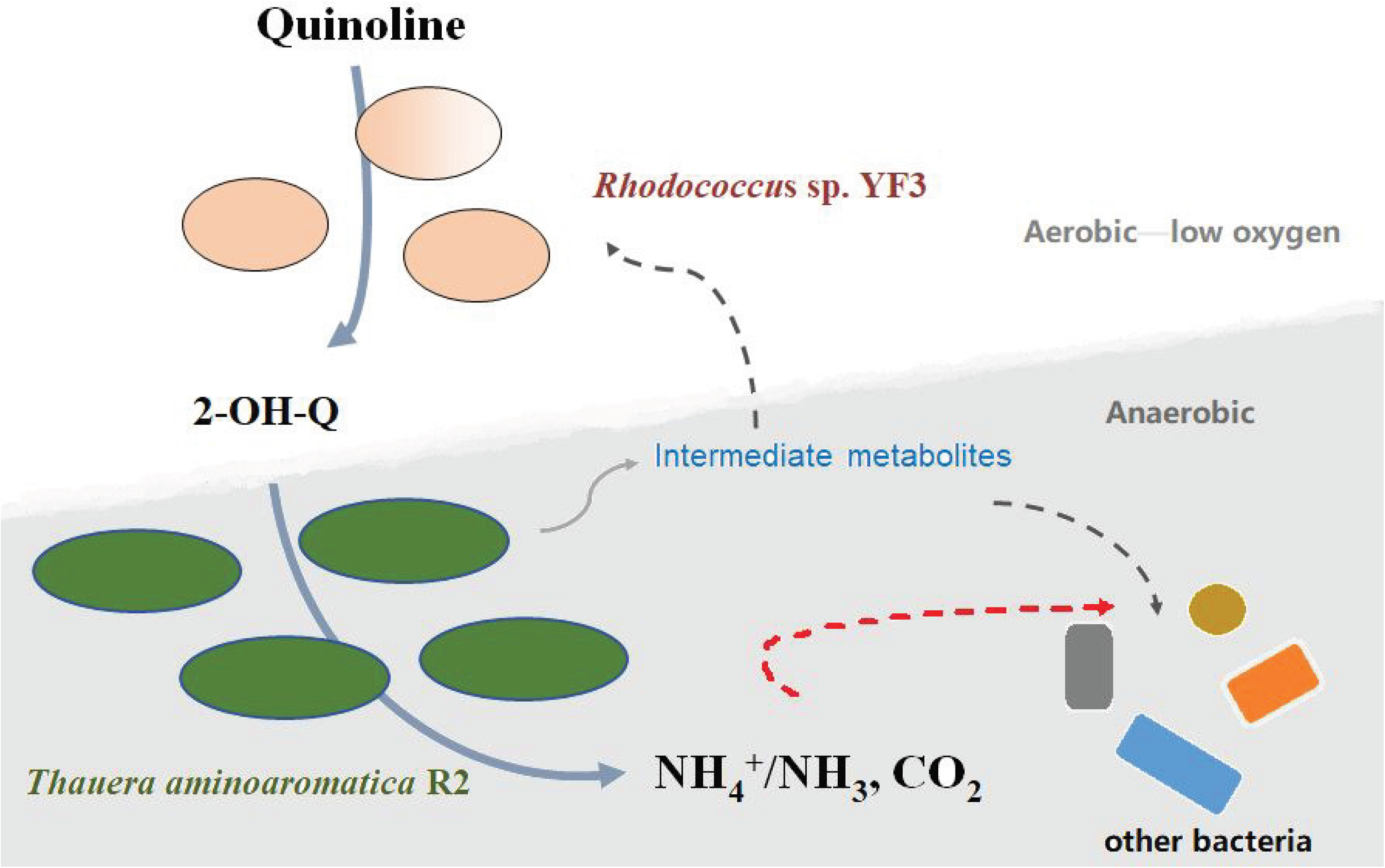
Proposed model for a syntrophic degradation of quinoline by *Rhodococcus* sp. YF3 and *Thauera* sp. R2.

Moreover, it was worth considering why this multi-strain degradation occurred in the quinoline-degrading reactor. In previous research (38), main reactions with respect to starch biodegradation occurred at the lower part of reactor while the small molecules such as acetate were converted to methane at the upper part. The organic compounds were gradually attenuated along the upflow direction. It was speculated that in our closed reactor, during the synthetic wastewater upflowing (i.e. from the bottom to up), the dissolved oxygen coming with the synthetic wastewater (ca. 7 mg/L dissolved oxygen) was rapidly consumed by *Rhodococcus* and other aerobic bacteria in the bottom of reactor, which then made an anoxic denitrifying niche in the inner space of bioreactor for *Thauera* to degrade 2-hydroxyquinoline that was produced by hydroxylation via *Rhodococcus* activity under limited oxygen or anoxic condition.

Metabolic cooperation between/among different bacterial members in the community are beneficial to the competitiveness. For instance, it has been shown that degradation of 4-chlorodibenzofuran by *Sphingomonas* sp. RW1 results in the accumulation of a dead-end product 3,5-dichlorosalicylate, while the inoculation with *Burkholderia* sp. JWS enables the cooperative complete degradation (39). A consortium comprised of *Escherichia coli* SD2 and *Pseudomonas putida* KT2440 pSB337 efficiently degrades parathion without accumulation of toxic intermediates (40). Besides, it has also been reported that four members were actively involved in the degradation of 4-chlorosalicylate, in which the strain of MT4 alleviated the stress of MT1 by transforming the toxic intermediate protoanemonin (41). These examples indicated that the metabolic complementarity in the microbial community were in favor of efficient bioremediation to contaminants and also benefited for the better adaptation or survival of bacteria in the specific environment.

This study collectively discussed commensal interaction in a quinoline-degrading consortium, aiming at highlighting importance of cross-feeding relationship between/among bacteria in biodegradation of organic compounds in the environment. Although our knowledge of this commensal relationship is not yet complete, understanding of such complex microbial interaction would provide useful information for assessing the biodegradability of organic compounds in the natural environment, for instance, the quinoline denitrifying degradation. Here we provided an unexpected and amazing insight into the microbial interaction. *Thauera aminoaromatica* R2, which adapted to denitrifying niche, could cooperate with *Rhodococc*us *pyridinivorans* YF3 to form a cross-feeding guild. To our best knowledge, this is the first report on cooperative relationship in quinoline denitrifying degrading guild, where two cross-feeding bacterial strains cooperated to degrade quinoline. However, the key bacteria, which open the food box for quinoline degradation, did not directly benefit from the hydroxylation under limited oxygen condition.

## Acknowledgment

This work was supported by the National Natural Science Foundation of China (NSFC 31670105 and NSFC 21177086).

